# *fusion*DB: assessing microbial diversity and environmental preferences via functional similarity networks

**DOI:** 10.1101/035923

**Authors:** C Zhu, Y Mahlich, M Miller, Y Bromberg

## Abstract

Microbial functional diversification is driven by environmental factors, *i.e.* microorganisms inhabiting the same environmental niche tend to be more functionally similar than those from different environments. In some cases, even closely phylogenetically related microbes differ more across environments than across taxa. While microbial similarities are often reported in terms of taxonomic relationships, no existing databases directly links microbial functions to the environment. We previously developed a method for comparing microbial functional similarities on the basis of proteins translated from the sequenced genomes. Here we describe *fusion*DB, a novel database that uses our functional data to represent 1,374 taxonomically distinct bacteria annotated with available metadata: habitat/niche, preferred temperature, and oxygen use. Each microbe is encoded as a set of functions represented by its proteome and individual microbes are connected via common functions. Users can search *fusion*DB via combinations of organism names and metadata. Moreover, the web interface allows mapping new microbial genomes to the functional spectrum of reference bacteria, rendering interactive similarity networks that highlight shared functionality. *fusion*DB provides a fast means of comparing microbes, identifying potential horizontal gene transfer events, and highlighting key environment-specific functionality.

*fusion*DB is publicly available at http://services.bromberglab.org/fusiondb/.

## Introduction

Microorganisms are capable of carrying out much of molecular functionality relevant to a range of human interests, including health, industrial production, and bioremediation. Experimental study of these microbes to optimize their uses is expensive and time-consuming; *e.g.* as many as three hundred biochemical/physiological tests only reflect 5-20% of the bacterial functional potential (Garrity GM, 2001). The recent drastic increase in the number of sequenced microbial genomes has facilitated access to microbial molecular functionality from the gene/protein sequence side, via databases like Pfam (Sayers, et al., 2009), COG (Tatusov, et al., 2003), TIGRfam (Haft, et al., 2003), RAST (Aziz, et al., 2008) and others. Note that the relatively low number of available experimental functional annotations limits the power of these databases in recognizing microbial proteins that provide novel functionality. Additional information about microbial environmental preferences can be found, *e.g.* in GOLD (Pagani, et al., 2012). While it is well known that environmental factors play an important role in microbial functionality (Cohan, 2001), none of the existing resources directly link environmental data to microbial function.

We mapped bacterial proteins to molecular functions and studied the functional relationships between bacteria in the light of their chosen habitats. We previously developed *fusion* (Zhu, et al., 2015), an organism functional similarity network, which can be used to broadly summarize the environmental factors driving microbial functional diversification. Here we describe *fusion*DB – a database relating bacterial *fusion* functional repertoires to the corresponding environmental niches. *fusion*DB is explorable via a web-interface by querying for combinations of organism names and environments. Users can also map new organism proteomes to the functional repertoires of the reference organisms in *fusion*DB; including, notably, matching proteins of yet unannotated function across organisms. The submitted organisms are visualized, and can be further explored, interactively as *fusion* networks in the context of selected reference genomes. Additionally, the web interface generates *fusion*+ networks, *i.e.* views that explicitly indicate shared microbial functions.

Our overall analyses of the *fusion*DB data for the first time give quantitative support for the fact that environmental factors driving microbial functional diversification. To demonstrate *fusionDB* functionality, for individual organisms we mapped a recently sequenced genome of a freshwater *Synechococcus* bacterium to *fusion*DB. In line with our previous findings (Zhu, et al., 2015), we demonstrate that this microorganism is more functionally related to other fresh water Cyanobacteria than to the marine *Synechococcus*. In a case study on *Bacillus* microbes we use *fusionDB* to track organism-unique functions and illustrate the detection of core-function repertoires that capture traces of environmentally driven horizontal gene transfer (HGT). *fusionDB* is a unique tool that provides an easy way of analysing the, often unannotated, molecular function spectrum of a given microbe. It further places this microbe into a context of other reference organisms and relates the identified microbial function to the preferred environmental conditions. Our approach allows for detection of microbial functional similarities, often mediated via horizontal gene transfer, that are difficult to recover via phylogenetic analysis. We note that *fusionDB* may also be useful for the analysis of functional potentials encoded in microbiome metagenomes. We expect that *fusionDB* will facilitate the study of environment-specific microbial molecular functionalities, leading to improved understanding of microbial lifestyles and to an increased number of applied bacterial uses.

## Methods

### Database setup

*fusion*DB is based on alignments of 4,284,540 proteins from 1,374 bacterial genomes (Dec. 2011 NCBI GenBank (Benson, et al., 2009). For each bacterium, we store its (1) NCBI taxonomic information (Sayers, et al., 2009) and, where available, (2) environmental metadata (temperature, oxygen requirements, and habitat; GOLD (Pagani, et al., 2012). The environments are generalized, *e.g. thermophiles* include hyper-thermophiles. “*No data*” is used to indicate missing annotations (Supplementary Online Material, SOM_Table 1). The general *fusion* (functional repertoire similarity-based organism network) protocol is described in (Zhu, et al., 2015). Briefly, all proteins in our database are aligned against each other using three iterations of PSI-BLAST (Altschul, et al., 1990) and the alignment length and sequence identity are used to compute HSSP (Rost, 2002). A network of protein similarities is then clustered using MCL (Dongen, 2000) clustering. For fusionDB the original *fusion* algorithm was modified to use less stringent protein functional similarity criteria (with HSSP distance cutoff = 10), which resulted in 457,576 functions (protein clusters; SOM Table 2). Each bacterium was thus mapped to a set of functions, its functional repertoire. Therefore our functional repertoires include all the bacterial functions, regardless of annotation. We are thus able to make function predictions, even the functions that have not been annotated before, for proteins in new bacteria.

### Web interface

*fusion*DB web interface has two functions: *explore* and *map new organisms*. The *explore* section contains access to all the 1,374 bacteria and their metadata. Users can search these with (combinations of) organism names and environmental preferences by using text box input or built in filters. User-selected organism set is then used to create a *fusion* network, in which organism nodes are connected by functional similarity edges. The *fusion* network can be viewed in an interactive display, as well as downloaded as network data files or static images. The user-defined color labels of the organism nodes reflect microbial taxonomy or environment. In the interactive display clicking an organism node reveals its taxonomic information and environmental preferences, while clicking an edge between two organisms yields a list of their shared functions. A *fusion*+ network can further be generated from the same list of organisms. There are two types of vertices (nodes) in *fusion+*: organism nodes and function nodes. Organism nodes are connected to each other <u>only</u> through the function nodes they share. The number of edges (degree) of an organism node represents the total number of functions of the organism; the relative position of each organism node is determined by the pull *towards* other organisms via the common functions and *away* from others via unique functions (Zhu, et al., 2015). Like *fusion*, *fusion+* can be interactively displayed, downloaded, and colored by the users’ choices. For both network types, users can further retrieve the functions shared by the selected organisms - the core-functional repertoire of the set. Note that the function annotation is from myRAST (including hypothetical function or unknown function; Aziz, et al., 2008). This feature is an efficient tool for investigating functions underlying organism diversification, particularly within different environment conditions.

In the *map* section, users can submit their own new organism proteomes (in fasta format) to our server. The submitted proteins are PSI-BLASTed against *fusion*DB and assigned to stored functions using the HSSP distance cutoff = 10. Note that novel proteins that can’t be assigned to existing functional groups are reported as functional singletons. Additionally, protein alignments that exceed 12 CPU hours of run-time are eliminated from future consideration. In testing, we found that no more than 0.1% of the proteins fall into this category. Although long run-times usually indicate that query proteins likely align to many others in our database, they contribute only a small fraction to the overall bacterial similarity and are eliminated for the sake of a faster result turn-around. The server sends out emails to users when mapping is finished. The *map* result page contains two tables: one is the list of functions of the submitted bacterium, while the other contains pairwise functional similarities (Eqn. 1) between the submitted bacterium and the reference proteomes in *fusion*DB (SOM Figure 1).

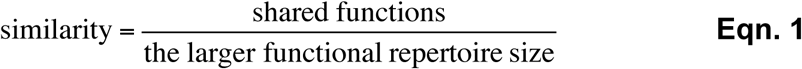

**Figure 1.**
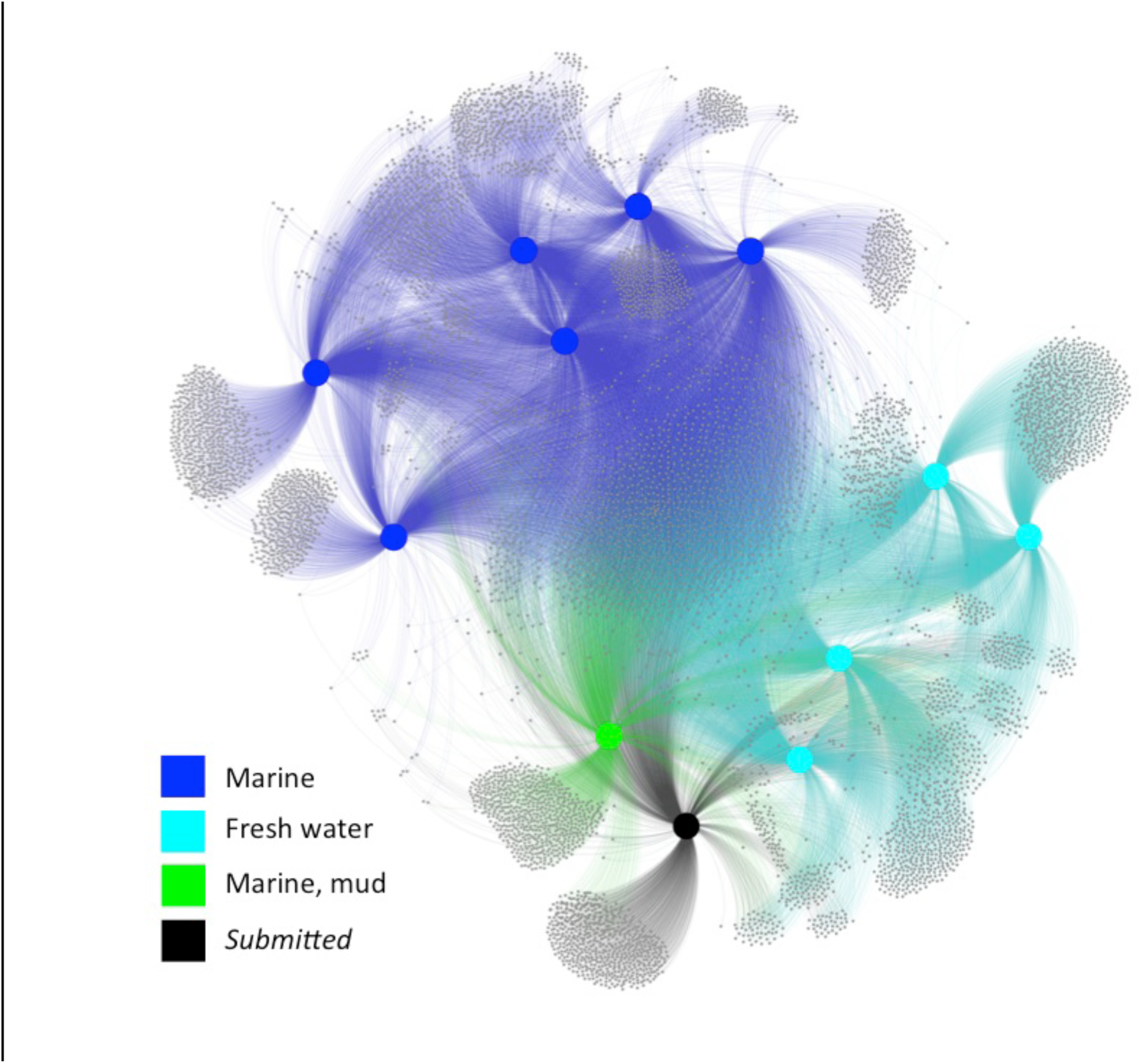
The fusion+ view of all Synechoccocus genomes. The submitted *Synechococcus sp.* PCC 7502 (black) cluster with the fresh water *Synechococcus* organisms (light blue). Note that the *Synechococcus sp.* PCC 7002 (green), which is isolated from marine mud, is salt tolerant but does not require salt for growth (see (Zhu, et al., 2015)).

Both tables can be easily sorted, searched and exported as comma-separated files. The submitted proteome is further mapped to user-selected reference organisms with *fusion* and/or *fusion*+ as describe above.

### Analysis of environment-driven organism similarity

For each environmental condition in *fusionDB*, we sampled organism pairs where organisms were from (1) the same condition (SC, *e.g.* both mesophiles) and (2) different conditions (DC, *e.g.* thermophile *vs*. mesophile). To alleviate the effects of data bias, the organisms in one pair were always selected from different taxonomic groups (different Families). The smallest available set of pairs, SC-psychrophile contained 33 organisms from 17 Families (SOM_Table 1; 136 pairs – 48 same phylum, 88 different phyla; due to high functional diversity of *Proteobacteria,* its classes were considered independent phyla). For all other environment factors we sampled, 100 bootstrap times, 136 organism pairs for both SC and DC sets, covering this same minimum taxonomic diversity. We calculated the pairwise functional similarity (Eqn. 1) distributions and discarded organism pairs with less than 5% similarity.

## Results and Discussion

### Map new Synechococcus genomes to fusionDB

We downloaded the full genome of *Synechococcus sp.* PCC 7502 (GCA_000317085.1) as translated protein sequence fasta (.faa file) from the NCBI Genbank (Benson, et al., 2009) and submitted it to our web interface. This 3,318 protein fresh water Cyanobacteria is isolated from a Sphagnum (peat moss) bog (Pagani, et al., 2012). 2,889 (87%) of the bacterial proteins mapped to 2,206 *fusion*DB functions and 426 (13%) were functional singletons; three proteins exceeded runtime and were excluded, Methods). The whole process from submission to receiving a results notification e-mail took a little under three and a half hours. The mapping indicates that *Synechococcus sp.* PCC 7502 is functionally most similar (56%) to *Synechocystis* PCC 6803, a fresh water organism closely related to *Synechococcus*. It also shares a high functional similarity with a mud *Synechococcus* (*S.sp.* PCC 7002; 53%) and with other fresh water *Synechococcus* (*S.elongatus* PCC 7942 and *S.elongatus* PCC 6301; 52%). Notably, but not surprisingly, *Synechococcus sp.* PCC 7502 shares much less functional similarity (40-42%) with the marine *Synechococcus* bacteria. This relationship is clearly demonstrated by the *fusion*+ networks (Figure 1). There are 874 functions shared by all the twelve *Synechococcus* (SOM Table 3), the core-function repertoire for this genus, and 1,128 functions shared among the fresh water *Synechococcus* (SOM Table 4). These additional 254 functions (SOM Table 5) are likely important for surviving in the fresh water, as opposed to the marine, environment, *e.g.* low salinity and low osmotic pressure.

### Environment significantly affects microbial function

Not surprisingly, the SC-thermophile and SC-psychrophile pairs demonstrate significantly higher similarities comparing to all DC pairs (Figure 2A). Notably, the higher functional similarity between thermophiles than between psychrophiles suggests that protein functional adaptation to low temperature is less drastic than to high temperature – an interesting finding itself. Contrast to the extremophiles, mesophile organisms seem to have huge functional diversity as the SC-mesophile similarities are comparable to those the DC pairs (Figure 2A).

**Figure 2.**
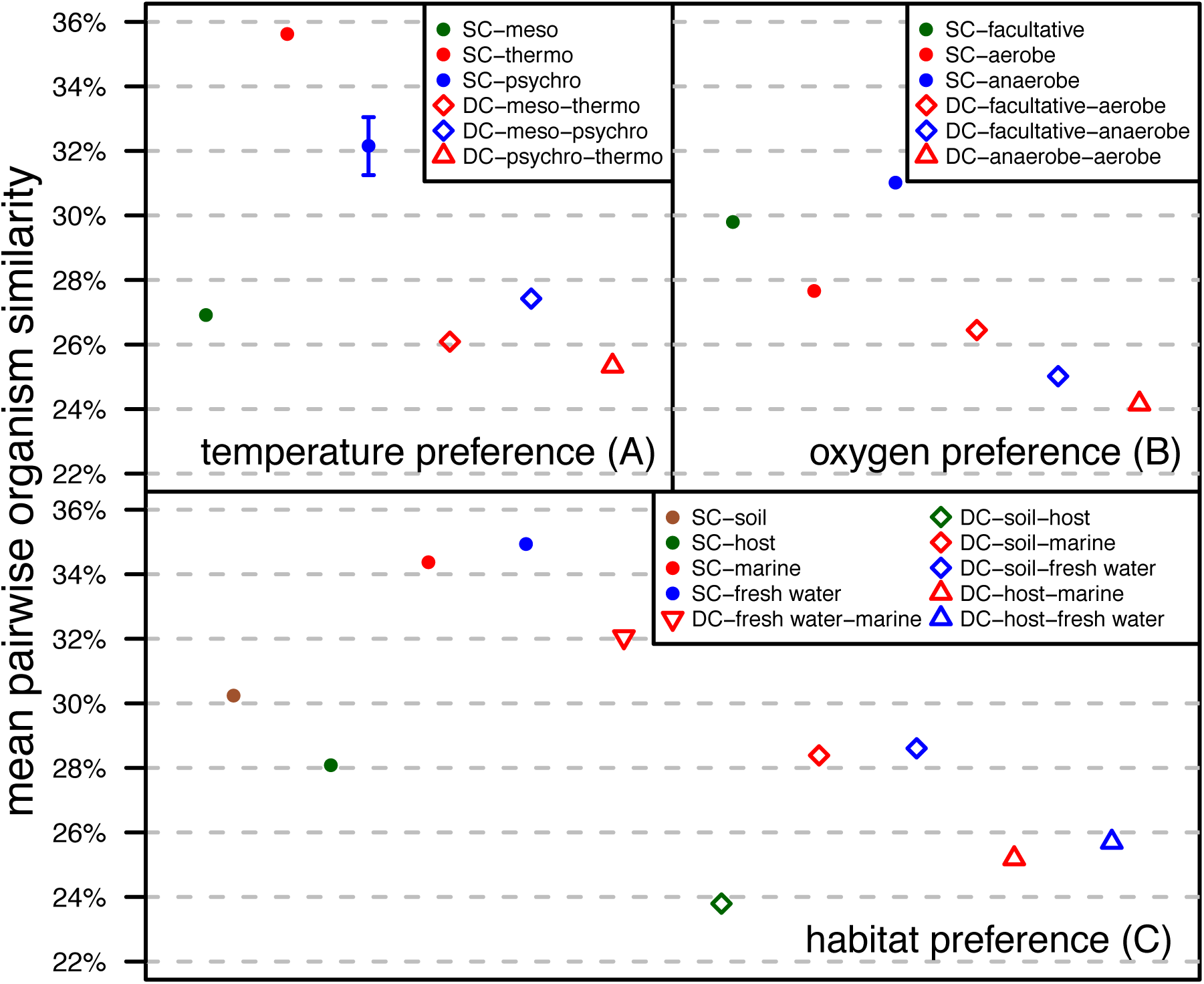
Organism pairwise similarity is higher among organisms living in the same environmental conditions. The mean pairwise similarity for same (SC) and different (DC) condition organisms according to (A) temperature preference, (B) oxygen requirement, and (C) habitat. For all points without error bars, the standard errors are vanishingly small.

Different molecular pathways of aerobic-respiration and anaerobic-respiration/fermentation explain the highest dissimilarity between the aerobes and anaerobes (DC-anaerobe-aerobe; Figure 2B). Interestingly, the SC-anaerobe similarities are higher than the SC-aerobe similarities, probably because the more ancient anaerobic-respiration/fermentation machinery is more simple and conserved.

Habitat-based DC samples show lower pairwise organism similarity than SC samples as well (Figure 2C), except for DC-fresh water-marine, which is not surprising due the same aquatic condition. SC-host displays the lowest mean organism similarity of the habitat SC samples. We speculate it is the result from the evolutionary pressure to deal with diverse host defence mechanisms (Hornef, et al., 2002). The soil organisms also share low functional similarity, which is likely due to soil’s heterogeneity at physical, chemical, and biological levels, from nano- to landscape scale (Bastian, et al., 2009).

In general, SC organisms across all environmental factors are more functionally similar than DC organisms (Figure 2; with exceptions mentioned above; Kolmogorov-Smirnov test p-val<2.5e-6). In other words, organisms in the same environment are generally more similar than organisms from different environments. This finding is intuitive and many studies have shown HGT within environment-specific microbiomes (Kim, et al., 2012; Liu, et al., 2012; Saye, et al., 1987). Our results, however, for the first time quantify on a broad scale the environmental impact on microorganism function diversification.

### Case study of a temperature driven HGT event

In *fusion*DB *explore*, we extracted thermophilic, mesophilic, and psychrophilic species representatives (one per species) of the *Bacillus* genus from *fusion*DB. We also added two other thermophilic organisms, *D. carboxydivorans* CO-1-SRB and *S. acidophilus* TPY, to generate a *fusion*+ network (SOM_Table 6, SOM_Figure 2). The non-*Bacillus* thermophiles were more closely related to the thermophilic *Bacilli*. All five thermophiles exclusively share three functions. One is a likely pyruvate phosphate dikinase (PPDK) that, in extremophiles, works as a primary glycolysis enzyme (Chastain, et al., 2011). Phylogenetic analysis (SOM Methods) suggests an HGT event between thermophilic organisms or a differential gene-loss in Bacilli that no longer live under high temperature (SOM Figure 3). The other two shared functions are carried out by proteins translated from mobile genetic elements (MGEs) that mediate the movement of DNA within genomes or between bacteria (Frost, et al., 2005). Shared closely related MGEs in distant organisms imply HGT (Krupovic, et al., 2013). We thus suggest that *fusion*DB offers a fast and easy way to trace functionally-necessary HGT within niche-specific microbial communities.

We have highlighted the importance of environmental factors for microbial function, and demonstrated the capability of *fusion*DB to not only annotate functions, but also directly link function to environment. Although it was developed for mapping new microbial genomes, *fusion*DB also has the potential for microbiome annotation. By mapping the proteins translated from metagenomes assembly to *fusion*DB, both the functional and taxonomical can be obtained. We look forward to making fusionDB more useful in this direction.

## 4. Conclusions

*fusion*DB links microbial functional similarities and environmental preferences. Our data analysis reveals environmental factors driving microbial functional diversification Mapping new genomes to the reference genomes, it offers a novel, fast, and simple way to detect core-function repertoires, unique functions, as well as traces of HGT. With more microbial genome sequencing and further manual curation of environmental metadata, we expect that *fusion*DB will become an integral part of microbial functional analysis protocols in the near future.

## Acknowledgements

We thank Drs. Burkhard Rost (TU Munich), Max Haggblom and Tamar Barkay (both Rutgers), and Tom O. Delmont (U Chicago) for all discussions and to those who deposit their data in public databases.

### Funding

This work was supported by the NSF CAREER Award 1553289, USDA-NIFA 1015:0228906 and the TU München – Institute for Advanced Study Hans Fischer Fellowship, funded by the German Excellence Initiative and the EU Seventh Framework Programme, grant agreement 291763.

### Conflict of Interest

none declared

